# A structure-function substrate of memory for spatial configurations in medial and lateral temporal cortices

**DOI:** 10.1101/2020.10.14.338947

**Authors:** Shahin Tavakol, Qiongling Li, Jessica Royer, Reinder Vos de Wael, Sara Larivière, Alex Lowe, Casey Paquola, Elizabeth Jefferies, Tom Hartley, Andrea Bernasconi, Neda Bernasconi, Jonathan Smallwood, Veronique Bohbot, Lorenzo Caciagli, Boris Bernhardt

**Author notes:** joint co-authors (Lorenzo Caciagli, Boris Bernhardt). **Corresponding Author:** Boris Bernhardt, PhD, Multimodal Imaging and Connectome Analysis Lab, McConnell Brain Imaging Centre, Montreal Neurological Institute and Hospital, McGill University, Montreal, Quebec, Canada.

## Abstract

Prior research has shown that structures of the mesiotemporal lobe, particularly the hippocampal-parahippocampal complex, are engaged in different forms of spatial cognition. Here, we developed a new paradigm, the Conformational Shift Spatial task (CSST), which examines the ability to encode and retrieve spatial relations between three unrelated items. This task is short, uses symbolic cues, and incorporates two difficulty levels and can be administered inside and outside the scanner. A cohort of 48 healthy young adults underwent the CSST, together with a set of validated behavioral measures and multimodal magnetic resonance imaging (MRI). Interindividual differences in CSST performance correlated with scores on an established spatial memory paradigm, but neither with episodic memory nor pattern separation performance, highlighting the specificity of the new measure. Analyzing high resolution structural MRI data, individuals with better spatial memory showed thicker medial as well as lateral temporal cortices. Functional relevance of these findings was supported by task-based functional MRI analysis in the same participants and *ad hoc* meta-analysis. Exploratory resting-state functional MRI analyses centered on clusters of morphological effects revealed additional modulation of intrinsic network integration, particularly between lateral and medial temporal structures. Our work presents a novel spatial memory paradigm and supports an integrated structure-function substrate in the human temporal lobe. Task paradigms are programmed in python and made open access.

## INTRODUCTION

Spatial memory is characterized by the encoding and retrieval of spatial associations. In rodents, structures of the mesiotemporal lobe (MTL) have long been recognized as crucial neural substrates of spatial memory (O’Keefe & Dostrovsky, 1971; O’Keefe & Nadel, 1978; Winocur, 1982; Morris, et al., 1982; Aggleton, Hunt, & Rawlins, 1986; Hafting, et al., 2005). In humans, early studies in patients with temporal lobe epilepsy revealed a direct correlation between the severity of MTL lesions and deficits in spatial cognition (Milner, 1965; Smith & Milner, 1981; Smith & Milner, 1989; Rains & Milner, 1994). Ensuing neuroimaging and lesion experiments in neurological patients reinforced the significance of the MTL as critical brain structures in spatial memory processing, but also pointed to an involvement of other brain regions and the broader conceptualization of spatial memory as a network-based phenomenon (Aguirre, et al., 1996; Ghaem, et al., 1997; Maguire, et al., 1998). The role of the MTL as a spatial processing hub was further supported by the discovery of human place cells and grid cells, specialized neurons believed to instantiate a scalable and navigable mental representation of space (Ekstrom, et al., 2003; Jacobs, et al., 2013).

The structural organization of spatial memory relies on the interplay between brain morphology and relevant cognitive phenotypes. For instance, the association between the volume of the MTL and behavioral measures of spatial cognition have been reported since the earliest structural magnetic resonance imaging (sMRI) studies (Abrahams, et al., 1999; Maguire, et al, 2000; Hartley & Harlow, 2012). Today, state-of-the-art automated segmentation tools can generate surface-wide representations of the brain, sampling morphological markers such as neocortical thickness and volume of hippocampal subregions with unprecedented resolution (Kim, et al., 2005; Fischl, FreeSurfer., 2012; Wang, et al., 2013; Caldairou, et al., 2016; Romero, Coupé, & Manjón, 2017; Goubran, et al., 2019). These millimetric anatomical indices are ideal for investigating the link between morphological and behavioral variability across individuals. Complementing sMRI studies, a large body of research has focused on the analysis of functional MRI (fMRI) acquisitions. Task-based fMRI studies have shown consistent MTL involvement during spatial memory tasks, together with activations in neocortical areas (Aguirre & D’Esposito, 1997; Jokeit, Okujava, & Woermann, 2001; Hassabis & Maguire, 2009; Schindler & Bartels, 2013). Complementing these paradigms, resting-state fMRI (rs-fMRI) enables to interrogate intrinsic functional networks (Biswal, Van Kylen, & Hyde, 1997; Cordes, et al., 2000; Lowe, Dzemidzic, Lurito, Mathews, & Phillips, 2000; Fox, et al., 2006; Smith, et al., 2009). An increasing body of rs-fMRI studies has also assessed intrinsic functional network substrates underlying interindividual differences in cognitive capacities (Smith, et al., 2015; Medea, et al., 2016; Sormaz, et al., 2017; He, et al., 2019).

The current study devised a new and open-access paradigm to assess spatial memory in humans and to elucidate the functional anatomy of spatial memory processing via structural and functional MRI analyses. We developed the Conformational Shift Spatial Task (CSST), a short, easy-to-use assessement tapping into the capacity to encode and retrieve spatial interdependencies between three conceptually unrelated objects. We administered the CSST to 48 healthy indivuals inside a 3T Siemens Magnetom Prisma scanner as part of a broader fMRI battery, which included additional testing probes for semantic memory, episodic memory, and mnemonic discrimination. Together with the semantic and episodic memory tasks, the CSST constitutes an integral part of a relational memory fMRI battery that can address structural and functional convergence and divergence across relational mnemonic domains. All three tests were homogenized by (i) implementing comparable visual stimuli, (ii) incorporating task difficulty modulation across two conditions (i.e., 28 easy trials & 28 difficult trials), (iii) using a 3-alternative forced choice trial-by-trial paradigm. Given that these tasks are designed to probe the different domains of relational memory, we hypothesized that behavioral scores on the CSST would correlate with performances on the semantic and episodic association tasks, with greater association observed between spatial and semantic domains (Moscovitch & Nadel, 1997; Moscovitch, et al., 2005; McNaughton, et al., 2006; Constantinescu, O’Reilly, & Behrens, 2016; Bellmund, et al., 2018; Mok & Bradley, 2019). We also evaluated participants on supplementary assessment tools outside the scanner, including the Four Mountains Task (FMT), an established spatial memory paradigm (Hartely, et al., 2007). We futher hypothesized that CSST performance would show strongest correlations with performance on the FMT as both tasks are devised to tap into the same relational domain, that is, spatial processing. In addition to its task-based portion, our protocol encompassed structural MRI as well as rs-fMRI acquisitions. We used these to assess associations between spatial memory scores and variations in MRI-derived morphological measurements of neocortical thickness and hippocampal volume across participiants. Although our surface-based analyses were regionally unconstrained, based on prior literature in humans and animals studying spatial memory (O’Keefe & Nadel, 1978; Morris, et al., 1982; Smith & Milner, 1989; Rains & Milner, 1994; Aguirre & D’Esposito, 1997; Abrahams, et al., 1999; Maguire, et al, 2000; Jokeit, Okujava, & Woermann, 2001; Hafting, et al., 2005; Hassabis & Maguire, 2009; Hartley & Harlow, 2012; Schindler & Bartels, 2013), we expected to observe structure-function substrates in medial temporal lobe regions. Results were contextualized against task-based fMRI findings in the same participants and *ad hoc* meta-analytical inference. Structural imaging observations were further used for *post hoc* explorations of rs-fMRI connectivity modulations by interindividual differnces in task performance.

## METHODS

### Participants

A total of 48 healthy adults (16 women, mean age ± SD = 29.71 ± 6.55 years, range: 19 to 44 years), recruited in 2018 and 2019, participated in our study and had normal or corrected-to-normal vision. Control participants did not have any neurological or psychiatric diagnosis. Our study was approved by the Research Ethics Committee of McGill University and participants gave written and informed consent upon arrival at the Montreal Neurological Institute.

### Conformational Shift Spatial Task

In the CSST, the participant discriminated the spatial arrangement of three semantically unrelated items (*i.e*., a brick, a tire, a bucket) from two additional foil configurations of the same items **(Figure 1a)**. At each trial, following a jittered inter-trial interval (1.5-2.5 seconds), the participant encoded the salient features of an original trio arrangement for a duration of 4 seconds. Following a jittered inter-stimulus interval (0.5-1.5 seconds), three distinct versions of the trio were displayed. All three conformations had undergone an equal rotation about the trio center of mass between 45° clockwise to 45° counterclockwise. The correct conformation had not undergone any additional transformation unlike the other two foils.

**Figure 1.**
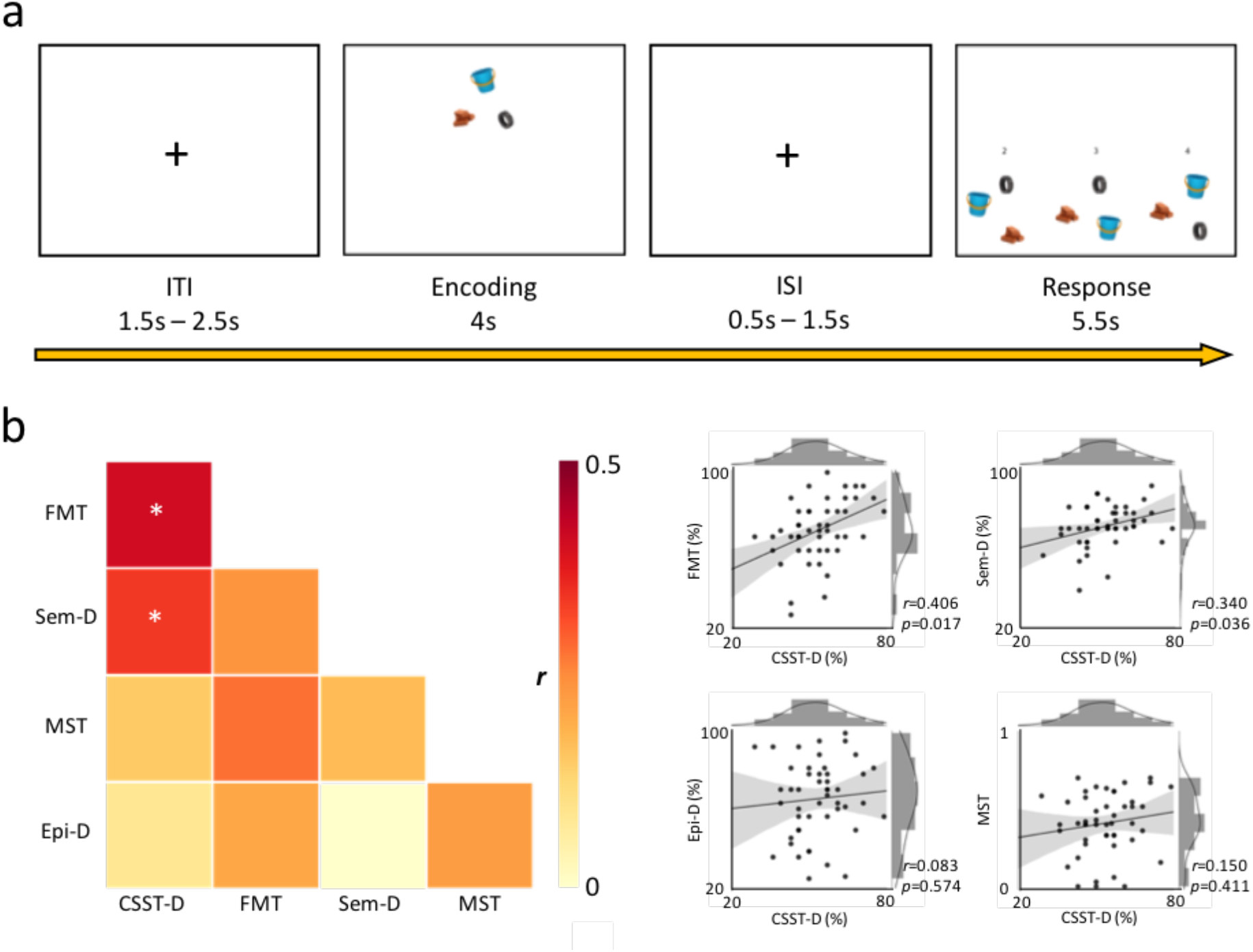
Task design and behavioral associations. **a)** Following a jittered inter-trial interval (ITI), the participant had to encode the spatial configuration of tire stimulus trio for 4 seconds. After a jittered inter-stimulus interval (ISI), the participant had 5.5 seconds to choose the original spatial conformation among two foil options, **b) left panel:** correlation heat map of performance across all tasks. CSST-D shows significant associations with FMT and Sem-D following adjustment for false discovery rate (\**pFDR*<0.05). **right panel:** joint-plot of CSST-D associations with other tasks (CSST-D: Conformational Shift Spatial Task-Difficult, FMT: Four Mountains Task, Sem-D: Semantic Task-Difficult, MST: Mnemonic Similarity/Discrimination Task, Epi-D: Episodic Difficult)

In the *difficult* condition (*i.e*., CSST-D), the two distractor layouts had been subjected to *one* specific additional transformation each: the spacing between the three items had changed with respect to the original configuration. In the *easy* condition (*i.e*., CSST-E), the foil configurations had undergone *two* specific additional transformations each: (1) the spacing between the three items had changed with respect to the original configuration; (2) the relative positions of trio items had been swapped. The participant was allowed up to 5.5 s to select the correct response. Thus, distractors in *difficult* trials varied from the original configuration by 2 degrees of separation (*i.e*., rotation about the center of mass and spacing alteration), whereas distractors in *easy* trials comprised 3 degrees (*i.e*., rotation about the center of mass, spacing alteration, and item positional swap).

The entire task was composed of 56 pseudo-randomized trials (*i.e*., 28 easy, 28 difficult). The semantic inter-relatedness of the trio items was computed via the UMBC Phrase Similarity Service (Han, Kashyap, Finin, Mayfield, & Weese, 2013). Based on the frequency with which two nouns representing the presented visual symbols co-occur within the Refined Stanford WebBase Corpus, which contains 100 million web pages from over 50,000 websites, this algorithm computed a conceptual relatedness index. We implemented prototypical visual stimuli as proxies for selected lexical entries whose similarity indices were inferior to 0.3 (range: 0-1).

### Additional cognitive tasks

#### Four Mountains Task (Hartley, et al., 2007)

The FMT is an established spatial cognition paradigm. In this version, 15 trials were administered in total. At each trial, the participant had 10 seconds to encode the spatially relevant stimuli within a computer-rendered landscape comprised of four distinct mountains varying in shape and size. After the encoding phase, participants had to select the correct landscape in a 4-alternative-forced-choice paradigm. The correct answer corresponded to the originally encoded landscape albeit depicted from a different first-person perspective, whereas the three incorrect options showed renderings of four mountains with different characteristics and configurations. All choices were additionally modified along lighting, weather, and vegetation texture to control for visual matching strategies. There was no time limit, but participants were instructed to respond as quickly and as accurately as possible. Following each trial, the participant had to report how certain they were about their response (*i.e*., certain or uncertain).

#### Semantic Task

We used a symbolic variant of a previously used lexicon-based semantic association paradigm (Sormaz, et al., 2017; Wang, et al., 2018). Consisting of 56 pseudorandomized trials, the task implements a 3-alternative-forced-choice paradigm and is modulated for difficulty across conditions with equal number of trials (*i.e*., 28 Difficult: Sem-D; 28 Easy: Sem-E). At each trial, a target object appeared at the top of the monitor (*i.e*., apple) with three objects below (*i.e*., desk, banana, kettle). Participants had to select the bottom item that was conceptually the most similar to the target. The semantic relatedness of items was measured via the UMBC similarity index (see above description regarding CSST). In difficult trials, the correct response and the target shared an index greater or equal to 0.7, whereas the foils shared a similarity index between 0.3 and strictly smaller than 0.7 with the target. In easy trials, the indices were greater than or equal to 0.7 between correct response and target, and 0 to strictly smaller than 0.3 between any given foil and target.

#### Episodic Task

We used symbolic variant of a previously used lexicon-based paradigm (Sormaz, et al., 2017; Payne et al. 2012) that involves two phases. In the encoding phase, participants had to memorize pairs of images shown simultaneously. Each pair was corrected for conceptual relatedness using the UMBC similarity algorithm (see above) with an index strictly smaller than 0.3. The encoding phase was modulated for difficulty across conditions: some trials were shown only once throughout the session, whereas others were displayed twice to ensure more stable encoding. Following a 10-minute delay, the retrieval phase was administered. At each trial, participants had to identify the object that was originally paired with the target object from the encoding phase in a 3-alternative-forced-choice paradigm, similar to the one described in the Semantic Task. There were 56 pseudo-randomized trials in total with 28 corresponding to pairs of images encoded only once (*i.e*., Epi-D) and 28, to pairs of images encoded twice (*i.e*., Epi-E).

#### Mnemonic Similarity Task (Stark, et al., 2013)

The MST assessed pattern separation, which is the capacity to hold orthogonal cognitive representations of overlapping stimuli. It comprised two phases: encoding and recall, administered ~8 minutes apart. The encoding phase consisted of 64 trials in which the participant had to choose whether the displayed item belonged “indoors” or “outdoors”. The recall phase was based on a 3-alternative-forced-choice paradigm. At this stage, the participant had to select whether the presented item was an exact duplicate from the encoding phase (*i.e*., “old”), an inaccurate duplicate (*i.e*., “similar”), or an altogether novel stimulus (*i.e*., “new”). This phase consisted of 32 trials per condition for a total of 96 trials.

### MRI acquisition

MRI data were acquired on a 3T Siemens Magnetom Prisma-Fit with a 64-channel head coil. Two T1-weighted (T1w) scans with identical parameters were acquired with a 3D-MPRAGE sequence (0.8mm isotropic voxels, matrix=320×320, 224 sagittal slices, TR=2300ms, TE=3.14ms, TI=900ms, flip angle=9°, iPAT=2). Task and resting-state fMRI time series were acquired using a 2D echo planar imaging sequence (3.0mm isotropic voxels, matrix=80×80, 48 slices oriented to AC-PC-30 degrees, TR=600ms, TE=30ms, flip angle=50°, multiband factor=6). The CSST task was approximately 15 minutes long and presented via a back-projection system to the participants. During the 7 minute-long rs-fMRI scan, participants were instructed to fixate a cross presented in the center of the screen and to not think about anything.

### Structural MRI processing

#### a) Generation of neocortical surfaces

To generate models of the cortical surface and to measure cortical thickness, native T1w images were processed using FreeSurfer 6.0 (http://surfer.nmr.mgh.harvard.edu). Previous work has cross validated FreeSurfer with histological analysis (Rosas, et al., 2002; Cardinale, et al., 2014) and manual measurements (Kuperberg, et al., 2003). Processing steps have been described in detail elsewhere (Dale, Fischl, & Sereno, 1999; Fischl, Sereno, Tootell, & Dale, 1999). In short, the pipeline includes brain extraction, tissue segmentation, pial and white matter surface generation, and registration of individual cortical surfaces to the fsaverage template. This aligns cortical thickness measurement locations among participants, while minimizing geometric distortions. Cortical thickness was calculated as the closest distance from the grey/white matter boundary to the grey matter/cerebrospinal fluid boundary at each vertex. Thickness data underwent spatial smoothing using a surface-based diffusion kernel (FWHM=10mm). As in prior work (Valk, et al., 2016), data underwent manual quality control and potential correction for segmentation inaccuracies.

#### b) Hippocampal subfield surface mapping

We harnessed a validated approach for the segmentation of hippocampal subfields, generation of surfaces running through the core of each subfield, and surface-based “unfolding” of hippocampal features (Caldairou, et al., 2016; Bernhardt, et al., 2016; Vos de Wael, et al., 2018). In brief, each participant’s native-space T1w image underwent automated correction for intensity non-uniformity, intensity standardization, and linear registration to the MNI152 template. Images were subsequently processed using a multitemplate surface-patch algorithm (Caldairou, et al., 2016), which automatically segments the left and right hippocampal formation into subiculum, CA1-3, and CA4-DG. An open-access database of manual subfield segmentations and corresponding high resolution 3T MRI data (Kulaga-Yoskovitz, et al., 2015) was used for algorithm training. A Hamilton-Jacobi approach (Kim, et al., 2014) generated a medial surface sheet representation running along the central path of each subfield and surfaces were parameterized using a spherical harmonics framework with a point distribution model (Styner, et al., 2006). For each subfield surface vertex, we then calculated columnar volume as a marker of local grey matter (Kim, et al., 2014). During data analysis, vertexwise projections of hippocampal columnar volume underwent surface-wide smoothing (FWHM=10) using SurfStat for Matlab (MathWorks, R2019b).

### Functional MRI processing

a. Task-based fMRI data were preprocessed using SPM12 (https://www.fil.ion.ucl.ac.uk/spm/). Steps included image realignment, distortion correction using AP-PA blip pairs, structural and functional co-registration, as well as functional data normalization and spatial smoothing (FWHM=6mm). Of the originally acquired fMRI scans, data for 4 participants were omitted due to artifacts caused by field inhomogeneity. For the remaining participants (n=44), first-level mass-univariate analyses were performed by modelling all task regressors into the SPM design matrix, which included trial onsets and durations/reaction times for ITIs, encoding phases, ISIs, retrieval phases, and post-retrieval rest periods, in addition to 6 standard motion parameters as well as a constant term. Regressors were convolved with the built-in SPM canonical hemodynamic response function without temporal nor dispersion derivatives. Following mass-univariate model estimations, first-level contrast maps from weighted comparisons between retrieval and encoding (*i.e*., when the participant chooses a specific stimulus configuration *vs*. when the participant is passively encoding the original stimulus conformation) were used to generate a single group-level activation map, which was thresholded (*pFWE*=0.05) and mapped onto fsaverage template using FreeSurfer.
b. The rs-fMRI scans were preprocessed using a combination of FSL, available at https://fsl.fmrib.ox.ac.uk/fsl/fslwiki (Jenkinson, et al., 2012), and AFNI, available at https://afni.nimh.nih.gov/afni (Cox, 1996), and included: removal of the first 5 volumes from each time series to ensure magnetization equilibrium, distortion correction based on AP-PA blip pairs, reorientation, motion correction, skull stripping, grand mean scaling, and detrending. Prior to connectivity analysis, time series were statistically corrected for effects of head motion, white matter signal, and CSF signal. They were also band-pass filtered to be within 0.01 to 0.1 Hz. All participants had overall low head motion and mean frame-wise displace. Following rs-fMRI preprocessing in native space, a boundary-based registration technique (Greve & Fischl, 2009) mapped the functional time series to each participant’s structural scan and subsequently, to the neocortical and hippocampal surface models. Surface-based fMRI data also underwent spatial smoothing (FWHM=10mm).

### Statistical analysis

Analyses were performed using SurfStat for Matlab (MathWorks, R2019b) available at http://math.mcgill.ca/keith/surfstat (Worsley, et al., 2009).

#### A) Behavioral task correlation

To assess the sensitivity and specificity of the newly developed protocol for spatial cognition, we cross-correlated the CSST-D with the FMT, Sem-D, Epi-D, and MST. Given that all participants were high functioning healthy individuals, we only incorporated performance scores on the difficult conditions where applicable, which additionally precluded ceiling effects.

#### B) Cortical thickness analysis

Surface-wide linear models evaluated task score and cortical thickness association:

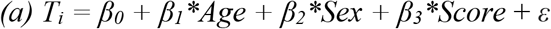

where *T_i_* is the thickness measure at vertex *i* for a total of 327,684 vertices. *Age, Sex*, and *Score* are model terms, *β_0_, β_1_, β_2_*, and*β_3_*, the estimated model parameters, and *ε* is the error coefficient. We then regressed out the effects of Age and Sex from cortical thickness measures:

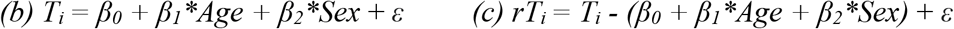

where *rT_i_* is the residual thickness measure at vertex *i*, corrected for *Age* and *Sex*. To assess whether the brain-behavioral correlations were generalizable to another spatial task, we correlated residual thickness from clusters of findings with FMT scores obtained outside the scanner.

#### C) Hippocampal analysis

Vertex-wise models also assessed effects of task scores on hippocampal columnar volumes:

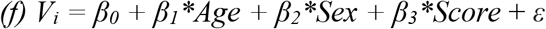

where *V_i_* is the local volume at vertex *i* for a total of 43,532 vertices. A similar model was also run for whole hippocampal volumes.

#### D) Functional contextualization

Task-based second-level functional activation maps were obtained from 44 participants and thresholded (*p*FWE=0.05) before being mapped to fsaverage. Average residual (*i.e*., age- and sex-corrected) cortical thickness across all vertices within regions of activation were then correlated with task scores. Furthermore, the meta-analytic platform of Neurosynth was used to perform a search for the term “navigation”, which resulted in 77 studies with a total of 3,908 activations. The generated association map was thresholded (*p*FDR=0.01) and mapped onto fsaverage. Once more, average residual thickness was computed and correlated with task results.

#### E) Resting-state connectivity analysis

Surface-wide linear models assessed the modulatory effect of task performance on rs-fMRI connectivity between clusters of structural imaging findings *(see B)* and resting-state data:

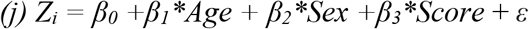

where *Z_i_* is the Fisher Z-transformed correlation coefficient between mean resting-state intensity for a given cluster in *(B)* and whole brain data at vertex *i*.

We performed a similar analysis to *(j)* to evaluate the effect of task score on functional connectivity between clusters in *(B)* and resting-state data mapped on the hippocampal template.

#### F) Correction for multiple comparisons

We used random field theory for non-isotropic images to correct for multiple comparisons (*p*FWE=0.05). Main structural MRI findings were based on a stringent cluster-defining threshold of *p*=0.001. For more exploratory rs-fMRI connectivity analyses, we used a more liberal cluster-defining threshold of *p*=0.025.

## RESULTS

### Behavioral f findings

We examined the association between the newly-developed conformational spatial shift task (CSST) and other tasks from our experimental protocol (**Figure 1b, Supplemental Table 1**). We excluded scores obtained on easy conditions across all difficulty-modulated tasks to prevent ceiling effects, as our cohort comprised of high functioning healthy adults (18.13±4.26 years of education; 47 currently employed/studying). Our participants indeed performed close to ceiling for the easy condition (CSST-E), but not the difficult condition (CSST-D) (*t*=16.8, *p*<0.001; **Supplemental Figure 1**). Furthermore, no sex differences were observed in CSST-D scores (**Supplemental Figure 2**). To ensure that the CSST is sensitive to spatial processing, we first cross-referenced it against the well-established four mountains task (FMT) paradigm that was administered outside the scanner (Hartley, et al., 2007). FMT scores correlated strongly with performances in both the CSST-E (*r*=0.419, *p*=0.003; **Supplemental Figure 3**) and the CSST-D (*r*=0.406, *p*=0.004; **Supplemental Figure 3**). Intra-CSST association was also significant (*r*=0.386; *p*=0.007; **Supplemental Figure 3**). CSST-D and FMT correlations were maintained when analyzing women and men separately (**Supplemental Figure 4**).

Several analyses supported specificity of CSST-D to spatial processing while also noting overlap with relational memory more generally (**Figure 1b**). Specifically, CSST-D also correlated with Sem-D (*r*=0.340; *p*=0.018) while showing neither an association with MST (*r*=0.150; *p*=0.308) nor with Epi-D (*r*=0.083; *p*=0.574). CSST-D also correlated to Sem-E (*r*=0.301; *p*=0.038), but not strongly to Epi-E (*r*=0.205, *p*=0.161; **Supplemental Figure 3**). As expected, CSST-D showed no meaningful associations with MST, Epi-D, and Epi-E when analyzing women and men separately, but only in men did CSST-D significantly correlate with Sem-D (**Supplemental Figure 4**).

### Structural substrates of spatial memory performance in neocortical regions

Controlling for age and sex, we observed positive correlations between CSST-D scores and thickness of bilateral superior temporal, left temporo-polar, bilateral parahippocampal, and left posterior cingulate cortices (**Figure 2a, Supplemental Figure 5**). Following correction for multiple comparisons (*p*FWE<0.05), findings were significant in the left superior temporal sulcus (*r*=0.597), left anteromedial superior temporal gyrus (*r*=0.609), right posterior parahippocampal gyrus (*r*=0.610), and the left inferior temporo-occipital junction (*r*=0.591; **Figure 2b**). CSST-D associations were consistent across clusters when separately analyzing both biological sexes (*r*values women/men; cluster 1: 0.59/0.62; cluster 2: 0.41/0.72; cluster 3: 0.66/0.62; cluster 4: 0.62/0.63; **Supplemental Figure 6**). Notably, average thickness of these four clusters also positively correlated with performance on the FMT (*r*=0.353; *p*=0.014; **Figure 2c**) and Sem-D (*r*=0.373; *p*=0.009; **Figure 2c**). Cluster-wise associations ranged between r=0.233-0.326 for FMT and between r=0.217-0.369 for Sem-D (**Supplemental Figure 7**). Although surface-based associations between thickness and 4MT were not significant after multiple comparisons correction, effect size maps were significantly similar to those from the correlation between thickness and CSST-D after correction for age and sex (r=0.472, non-parametric p<0.001) **(Supplemental Figure 8).** Cortical thickness did not correlate with scores in other tasks for the same significance criteria, indicating specificity of the observed brain-behavior correlations. These findings implicate local regions within the left temporal lobe as well as the right MTL as cortical substrates underlying interindividual differences in aptitude on the CSST-D.

**Figure 2.**
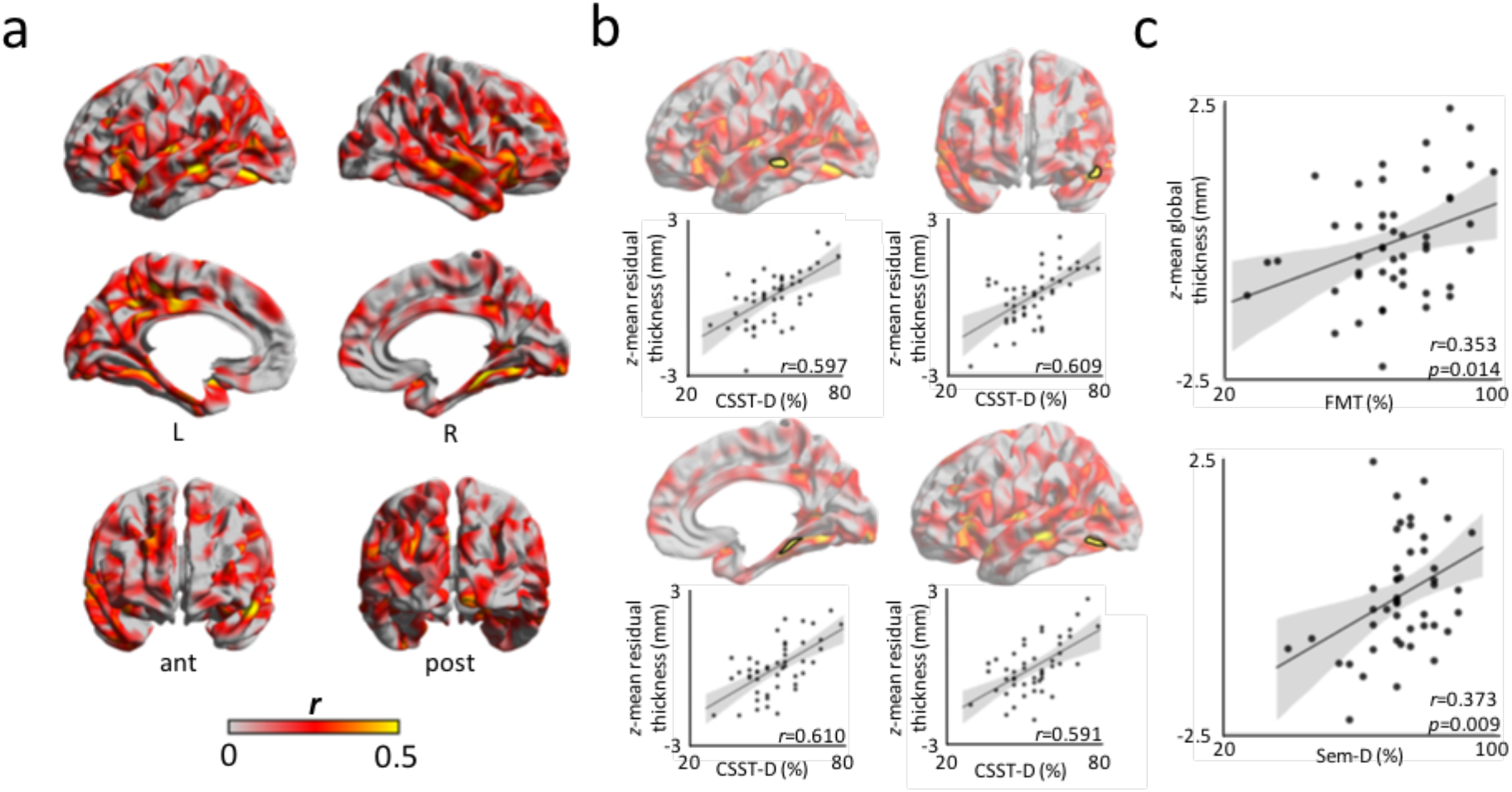
Cortical substrates of the CSST. **a)** Pearson correlation coefficients of CSST-D performance on neocortical thickness after regressing out age and sex. b) Findings shown after multiple comparisons correction(*p*FWE<0.05; cluster-defining threshold of CDT=0.001), highlighting clusters in the left superior temporal sulcus, left anteromedial superior temporal gyms, right posterior parahippocampal gyrus, and left inferior temporo-occipital junction, c) Correcting for age and sex, average cortical thickness across clusters of finding showed robust correlations with performance on FMT (*r*=0.353; *p*=0.014) and Sem-D (*r*=0.373; *p*=0.009).

### Structural substrates of spatial memory performance in hippocampal subregions

Controlling for effects of age and sex, we observed a strong trend between CSST-D scores and total hippocampal volume (r=0.234, one-tailed p=0.052). While no surface-wide association passed stringent criteria for multiple comparisons corrections (i.e., pFWE<0.05; CDT=0.001), we observed uncorrected associations between CSST-D and hippocampal columnar volumes along the long axis of each subfield **(Supplemental Figure 9)**.

### Functional contextualization

We contextualized the structural imaging findings with respect to areas relevant for spatial cognition, using task-based fMRI activation maps obtained from the same participants and Neurosynth-based meta-analysis. We pooled data across CSST-E and CSST-D trials (**Supplemental Table 2**) as one-tailed t-tests failed to ascertain significant group-level activation differences between conditions. We then mapped the volumetric second-level activations (**Supplemental Figure 10**, **Supplemental Table 3**) to fsaverage, and computed average cortical thickness in highlighted regions, which showed no correlation with CSST-D scores (*r*=0.189, one-tailed *p*=0.110; **Figure 3a**). An additional *ad hoc* meta-analysis was also performed (**Figure 3b**); here, the Neurosynth-derived map was similarly mapped to fsaverage and average cortical thickness in activated areas was computed. We observed a significant association between CSST-D behavioral performance and average thickness across Neurosynth-derived regions (*r*=0.319, one-tailed *p*=0.014).

**Figure 3.**
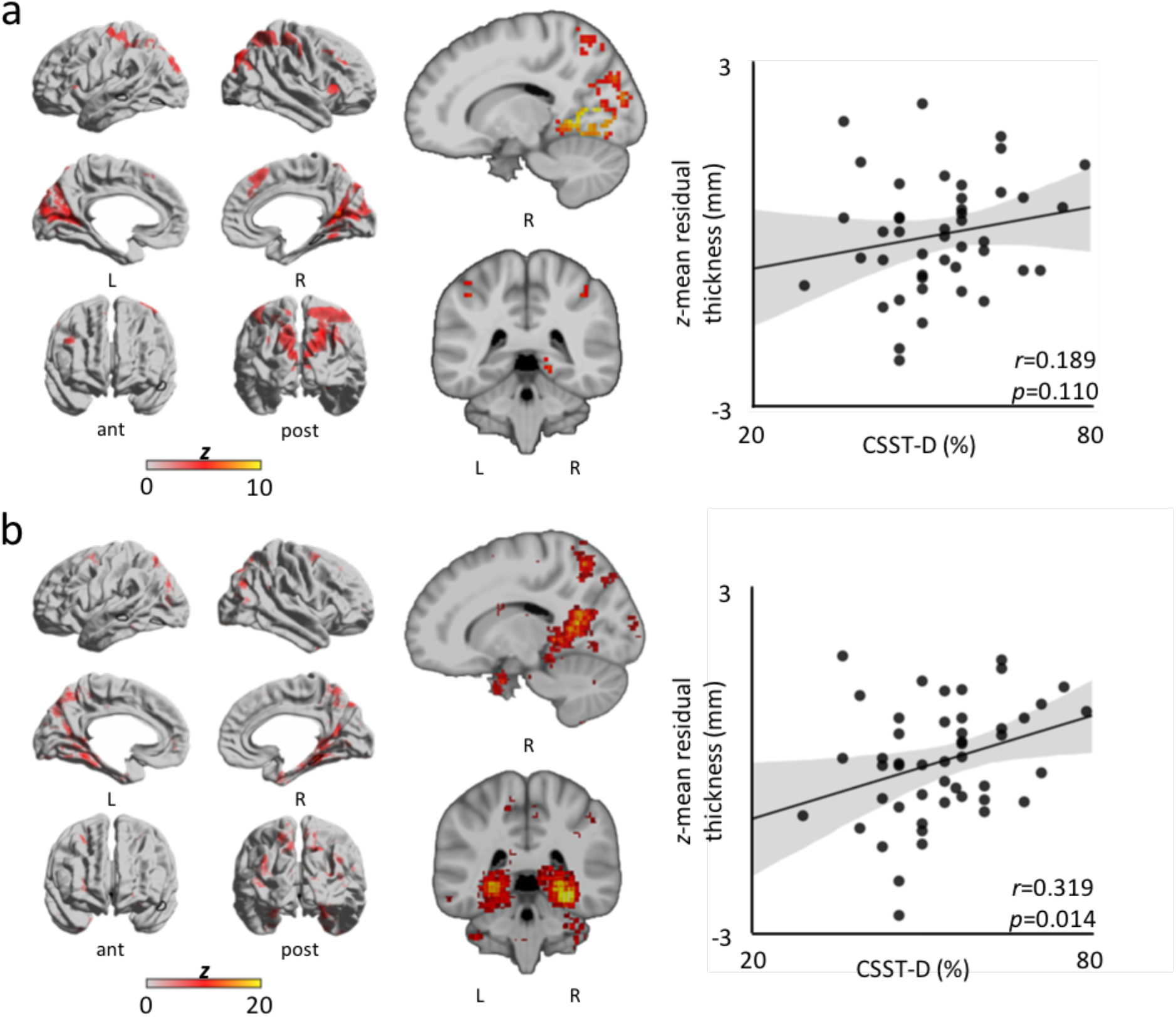
Functional contextualization. To further address task validity, CSST-D scores were correlated with average cortical thickness across regions of activation from CSST- and Neurosynth-derived maps, **a)** *Left column:* Group level (n=44) CSST surface-wide activation for *retrieval-vs.-encoding* weighted contrast. Notably, the right PHG shows significant activation. Dark outlines correspond to structural clusters. *Middle column:* volumetric activation (MNI coordinates: 15, −40). *Right column:* Trend between average cortical thickness across regions of activation and CSST-D score (r=0.189, one-tailed p=0.110). b) *Left column:* Neurosynth-derived surface-wide coactivations for the term *“navigation”*. Dark outlines correspond to structural clusters. *Middle column:* volumetric activation (MNI coordinates: 15, −40). *Right column:* Significant association between average cortical thickness across coactivated areas and CSST-D performance (*r*=0.319, one-tailed *p*=0.014).

### Modulatory effect of task performance on functional connectivity profile

We conducted exploratory seed-based connectivity analyses centered on clusters of findings from the structural analyses (*i.e*., left superior temporal sulcus, left anteromedial superior temporal gyrus, right posterior parahippocampal gyrus, and the left inferior temporo-occipital junction) **(Figure 4)**. Accounting for age and sex, we observed a marginal association between CSST-D score and the connectivity strength of the right parahippocampal cluster (seed 3; **Figure 4a**) and a region encompassed by the left middle frontal and precentral gyri extending medially via the paracentral lobule into the anterior cingulate (*p*FWE=0.052; outlined cortical surface on 3^rd^ row; **Figure 4b**). Here, individuals with higher scores on CSST-D presented with higher functional connectivity between these nodes. We also found that CSST-D performance positively modulated connectivity between the left superior temporal sulcus (seed 1; **Figure 4a**) and left CA1-3 (*p*FWE=0.014; outlined hippocampal surface on 1^st^ row; **Figure 4b**). A similar modulation was seen for the cluster in the left inferior temporo-occipital junction (seed 4; **Figure 4a**), which showed connectivity modulation to right CA1-3 by CSST-D (*p*FWE=0.036; outlined hippocampal surface on 4^th^ row; **Figure 4b**)

**Figure 4.**
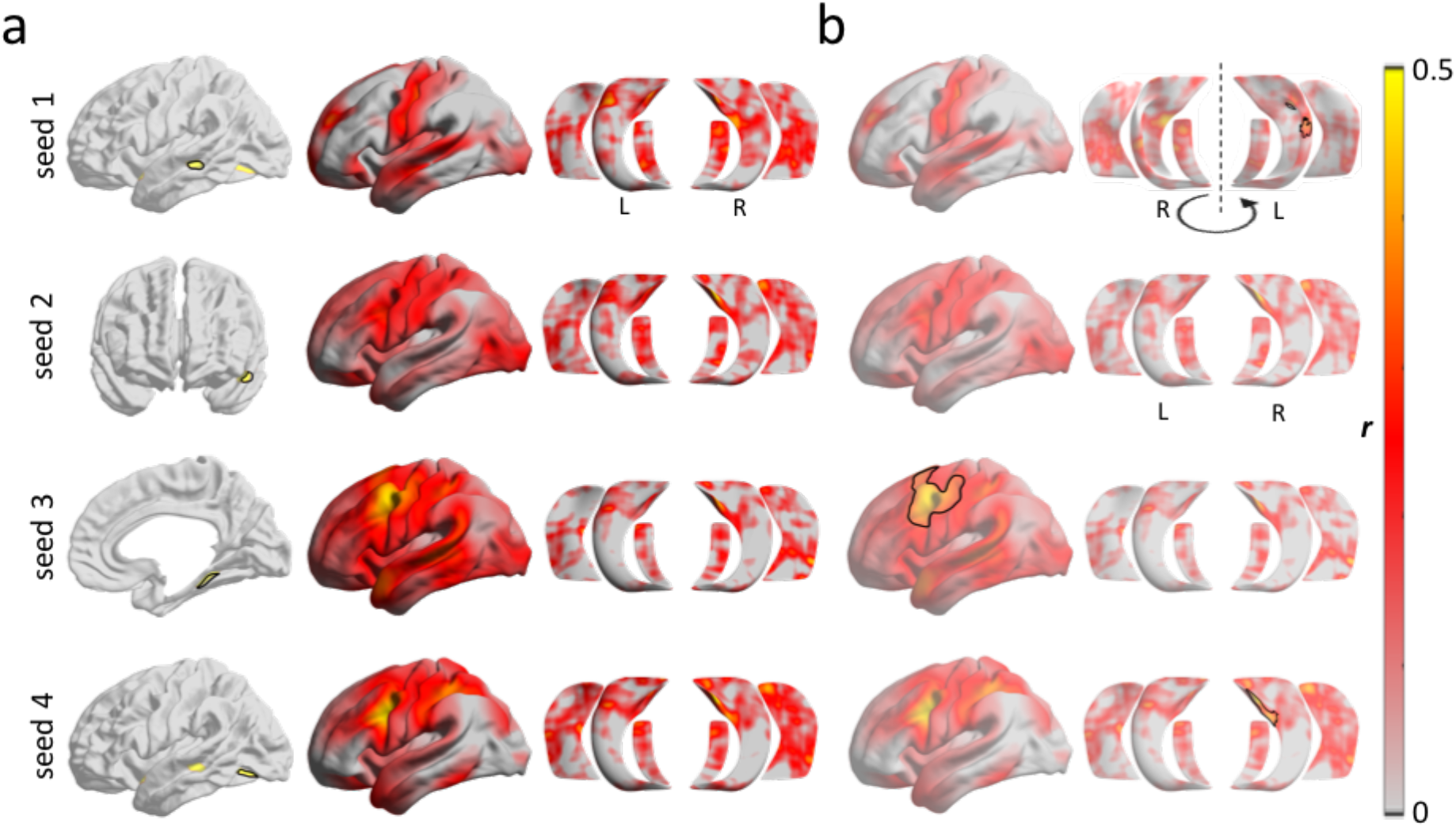
Seed-based resting-state functional connectivity analysis. Analyses was focussing on clusters of significant structural modulations (see *Figure 2*; presented in rows), **a)** *Left column:* seeds; *Middle & right columns*: whole-brain & hippocampal functional connectivity for each seed, b) Associations between CSST-D and connectivity profiles.

## DISCUSSION

Our goal was to design a novel neurocognitive task to evaluate the capacity for encoding and retrieving spatial relationships between unrelated objects in humans, and to identify brain substrates of such spatial processing via structural and functional connectivity analyses. To this end, we developed and administered the new Conformational Shift Spatial Task (CSST) to 48 healthy young adults as part of a broader task-based fMRI battery and conducted structural and resting-state fMRI (rs-fMRI) analyses. In addition to the CSST, our fMRI battery also included a semantic association task and an episodic memory task that were developed in consort with the CSST to address questions pertaining to relational memory more generally. All three tasks were homogenized in terms of visual stimuli, task difficulty and duration, as well as response paradigm (i.e., 3-alternative forced choice). These tests were further optimized for administration outside the scanner as well as inside, and are openly available. An additional test for assessing mnemonic discrimination was also included. Behavioral correlations with additional memory metrics supported relative sensitivity and specificity of the CSST to spatial memory, and some overlap with relational memory more generally. Studying *in vivo* measures of cortical morphology, we identified substrates underlying interindividual differences in CSST performance in a network of lateral and medial temporal lobe regions. Complementary explorations of rs-fMRI data indicated a stronger functional connectivity of these areas in individuals with higher scores on the CSST. Structural MRI findings could be functionally contextualized by showing overlaps to task-based fMRI activations from the CSST paradigm itself as well as *ad hoc* meta-analysis. In this work, we present a new paradigm that taps into spatial memory processing, and our multimodal MRI results offer new insights into integrated structure-function substrates of human spatial cognition.

The CSST is an openly accessible (https://github.com/MICA-MNI/micaopen) and convenient python-based protocol that can be administered inside or outside the scanner in less than fifteen minutes. It implements symbolic stimuli in a 3-alternative-forced-choice paradigm and consists of two experimental conditions modulated for difficulty (Easy: CSST-E; Difficult: CSST-D), which is suitable for the study of interindividual variations and between-group differences in the context of healthy and clinical cohorts. The CSST encompasses 56 pseudo-randomized trials (28 per condition) with four equivalent iterations, which can be leveraged to perform multiple probes while controlling for habituation. In addition to paradigm development, we assessed behavioral associations between CSST performance to measures obtained from tasks tapping into spatial, semantic, and episodic dimensions of memory. As this study analyzed high functioning healthy adults, we restricted the analyses to scores obtained on the difficult condition, CSST-D. The CSST-E scores, where our healthy individuals perform close to ceiling, may be more suitable for phenotyping individuals with deficits in spatial cognition, including older adults (Perlmutter, et al., 1981; Pezdek, 1983; Bohbot, et al., 2012) and those with neurological disorders (Bird, et al., 2010). In our cohort, CSST-D results correlated with FMT scores measured outside the scanner, suggesting that the task is sensitive to topographic memory. Interestingly, behavioral outcome on the CSST-D was neither correlated with scores on an episodic paired-associates task nor with performance on a pattern separation task. However, we did observe a correlation with a semantic decision making task, and in a prior study we had also found that the spatial and semantic aspects of memory were associated via the organization of connectivity between hippocampus and lateral temporo-parietal cortex (Sormaz, et al., 2017).

Following these behavioral explorations, we leveraged the CSST to determine potential structural correlates of interindividual differences in spatial cognition. We examined whether interindividual differences in CSST-D scores correlated to MRI-derived neocortical thickness and hippocampal columnar volume measures. Accounting for variance explained by age and sex, we observed associations with the thickness of bilateral superior temporal, left temporo-polar, bilateral parahippocampal, and left posterior cingulate areas. Following multiple comparisons correction, findings clustered within left lateral temporal and right medial temporal lobe areas, notably the right posterior parahippocampus. As a primary relay between the allo-cortical subregions of the hippocampal formation and the isocortex, the parahippocampal cortex plays an essential role in many different forms of spatial processing, including memory for scenes and configuration of objects (Aguirre, et al., 1996; Bohbot, et al., 1998; Epstein & Kanwisher 1998; Abrahams, et al., 1999; Bohbot, Allen, & Nadel, 2000; Bohbot et al., 2015). Increased gray matter volume of the entorhinal cortex has previously been associated with improved performance on games that rely on geometric relationships, such as Tetris and Minesweeper, as well as platform games, such as Super Mario 64 (Kühn & Gallinat, 2014). One study also found an increase in gray matter thickness of bilateral parahippocampal cortex following 15 daily gaming sessions on a first-person shooter platform, with long-lasting changes in the left parahippocampal cortex (Momi, et al., 2018). The authors argued that detailed environmental mapping of the virtual arena conferred a competitive advantage as evidenced by continued navigation during episodes of virtual blindness (*i.e*., when hit by smoke or flashbang grenades). However, too great a reliance on the response strategy mediated by the caudate nucleus, which is the most favored spontaneous navigational behavior in first-person shooter paradigms, has instead been shown to shrink the hippocampus (West, et al., 2018). Functional neuroimaging paradigms have further implicated the parahippocampal gyrus in object-location retrieval (Owen, et al., 1996), local geometry encoding (Epstein & Kanwisher, 1998; Epstein, 2008), fine-grained spatial judgment (Hirshhorn, et al., 2012), and 3D space representation (Kim & Maguire, 2018). In line with previous findings, our observations suggest that measures of parahippocampal gray matter could serve as a proxy for cortico-hippocampal information coherence, with greater efficiency of the system translating into better spatial cognition skills. Although their core microstructural changes are incompletely understood, it has been suggested that variations in cortical thickness may, nonetheless, capture underlying variations in cytoarchitecture. For example, while thickness measurements may be anti-correlated to neuronal density, regions of relatively high thickness with reduced density may instead present with more complex dendritic arborization, which could facilitate integrative information processing (Collins, et al., 2010; la Fougère, et al., 2011; Cahalane, Charvet, & Finlay, 2012; Wagstyl, et al., 2015). Regarding the hippocampus, we observed a positive trend between CSST-D scores and total volume (which were found to be distributed along the longitudinal axis). The hippocampus has long been associated with spatial processing in experimental work in animals (O’Keefe & Nadel, 1978; Aggleton, Hunt, & Rawlins, 1986; Sargolini, et al., 2006; Burgess, Barry, & O’Keefe, 2007) as well as patient lesion (Milner, 1965; Smith & Milner, 1981; Smith & Milner, 1989; Rains & Milner, 1994) and human neuroimaging studies (Aguirre, et al., 1996; Ghaem, et al., 1997; Maguire, et al., 1998; Abrahams, et al., 1999; Maguire, et al, 2000; Hassabis, et al., 2009; Robin, Buchsbaum, & Moscovitch, 2018; Kim & Maguire, 2018). Furthermore, task-based fMRI analysis of the CSST paradigm and ad hoc meta-analysis via Neurosynth confirmed consistent activations in the hippocampus-parahippocampus complex, particularly towards posterior divisions. It is worth noting that while our result pertaining to the overall volume of the hippocampus corroborates prior evidence, our analytical approach may not have been sensitive enough to identify subregional effects. Further analyses with larger cohorts and/or higher resolution imaging of the hippocampus are required to more robustly explore subregional substrates in the hippocampus; these approaches may benefit from approaches that tap into hippocampal longitudinal and medio-lateral axes (Vos de Wael, et al., 2018; Plachti, et al., 2019; Przeździk, et al., 2019; Paquola, et al., 2020).

In addition to results pertaining to the MTL, we observed structural MRI effects in lateral temporal areas whose role is less well defined in spatial cognition. Within-sample fMRI and *ad hoc* metaanalysis results did not support any functional relevance of lateral temporal regions to spatial memory processing and navigation. Several studies have already pointed to contributions from extra-MTL structures such as the posterior cingulate, retrosplenial, dorsolateral prefrontal, and posterior parietal cortices (Aguirre, et al., 1996; Ghaem, et al., 1997; Maguire, et al., 1998; Byrne, Becker, & Burgess, 2007; Whitlock, et al., 2008), but none explicitly to lateral temporal areas. In one virtual-reality fMRI study in which participants navigated a circular platform, grid-cell firing patterns were consistently observed in the lateral temporal cortices, in addition to MTL findings (Doeller, Barry, & Burgess, 2010). Another experiment showed that movement-onset periods in a square virtual environment were linked to increases in theta frequency power mainly within the hippocampus, but also across the lateral temporal lobes, with greater changes in theta power for relatively longer path lengths (Bush, et al., 2017). A meta-analytic review also found that bilateral lateral temporal cortices participated in the processing of familiar as opposed to recently learned virtual environments (Boccia, Nemmi, & Guariglia, 2014). Interestingly, the same review also reported greater involvement of the right parahippocampus in recently learned virtual settings when compared to familiar ones. Our observation that measurements of cortical thickness across disparate clusters within the left lateral temporal lobe correlate with performance on the CSST-D corroborates previous findings regarding a complementary role of lateral temporal areas to medial regions in spatial processing.

To provide a network-level context for these structural findings, we implemented seed-based rs-fMRI connectivity analyses centered on lateral and medial temporal clusters where morphological associations to CSST-D performance had been observed. Using more exploratory thresholding, we observed a positive association between CSST-D scores and the connectivity strength between components of this network, specifically between medial and lateral temporal regions, together with a region denoted laterally by the middle frontal and precentral gyri, and medially by the paracentral lobule and superior anterior cingulate. Significant connectivity modulations were obtained for three out of four clusters that showed main effects of CSST-D on cortical thickness; such a combined effect on morphology and functional connectivity speaks to intracortical and network level substrates underlying spatial cognition. Since the discovery of rodent place cells (O’Keefe & Dostrovsky, 1971) and the formulation of the cognitive map theory (O’Keefe & Nadel, 1978), which primarily focused on the hippocampus, the neural landscape of spatial cognition has increasingly been conceptualized as a network that encompasses widespread brain areas that perform complementary operations. For example, one leading model posits a vast circuit involving MTL and extra-MTL regions that participate in the reciprocal transformation of body-centered and subject-invariant spatial representations (Byrne, Becker, & Burgess, 2007; Dhindsa, et al., 2014; Bicanski, & Burgess, 2018). An overlooked assumption made by several previous models is that regions involved in specific neural processes may be recruited in various other cognitive domains. Given that the human brain is a finite organ capable of multiple mental functions, it is not surprising that many neural operations show anatomical convergence. In fact, some of the regions discussed herein in the context of spatial memory may apply equally as well to other related cognitive faculties (Constantinescu, O’Reilly, & Behrens, 2016; Epstein, et al., 2017; Bellmund, et al., 2018; Mok & Bradley, 2019).

Thus, the behavioral correlation that we observed between spatial and semantic memory scores could point to shared mechanisms across different mnemonic domains. This finding is in line with prior literature suggesting such functional versatility of the hippocampus, which is likely predicated on its structural connectivity to other brain systems (Moscovitch & Nadel, 1997; Moscovitch, et al., 2005). Notably, associations between semantic and spatial processing also paralleled our recent study of individual differences in different types of memory (Sormaz et al., 2017). In this study, we found that both semantic and spatial memory were related through their association between hippocampal and lateral parietal connectivity at rest. It has been proposed that the brain may organize semantic information as a navigable conceptual mental space, a mechanism not unlike the encoding of spatial information into a cognitive map via the consorted activity of hippocampal place cells and entorhinal grid cells (O’Keefe & Dostrovsky, 1971; Hafting, et al., 2005; McNaughton, et al., 2006; Constantinescu, O’Reilly, & Behrens, 2016). New evidence further indicates that these cell populations are in fact functionally more flexible than previously believed. For example, it has been argued that the neural mechanisms that encode for Euclidean space may also eventuate a multitude of orthogonally-stable cognitive spaces, each representing a unique dimension of experience, such as conceptual knowledge (Bellmund, et al., 2018). Recent findings support the involvement of domain-invariant learning algorithms that apply to the neural organization of both spatial and semantic information (Mok & Bradley, 2019). By implementing our newly-developed Conformational Shift Spatial Task in conjunction with stimulus-matched episodic and semantic memory paradigms, it may be possible to efficiently explore the degree of structural and functional convergence across relational memory domains both in healthy as well diseased populations.

