## Supplementary Materials for "A structure-function substrate of memory for spatial configurations in medial and lateral temporal cortices"

### SUPPLEMENTARY FIGURES

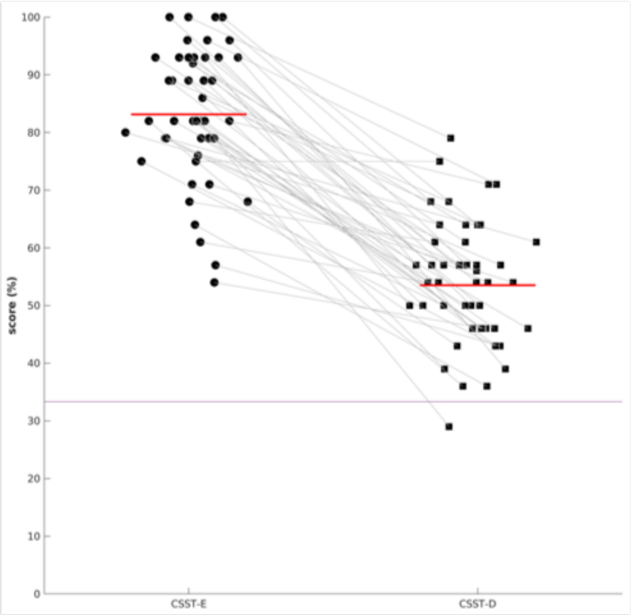


**Supplemental Figure 1 |** Participants scored significantly higher on the CSST-E (83.1±11.3%) compared to the CSST-D (53.5±10.7) as evidenced by a two-tailed paired student t-test ($t$=16.8, $p$<0.001). Red horizontal lines show distribution means. Chance level performance is depicted as a horizontal line (33.33%). Participants scored significantly higher than chance level on each condition (CSST-E: $t$=30.4, $p$<0.001; CSST-D: $t$=13.1, $p$<0.001).

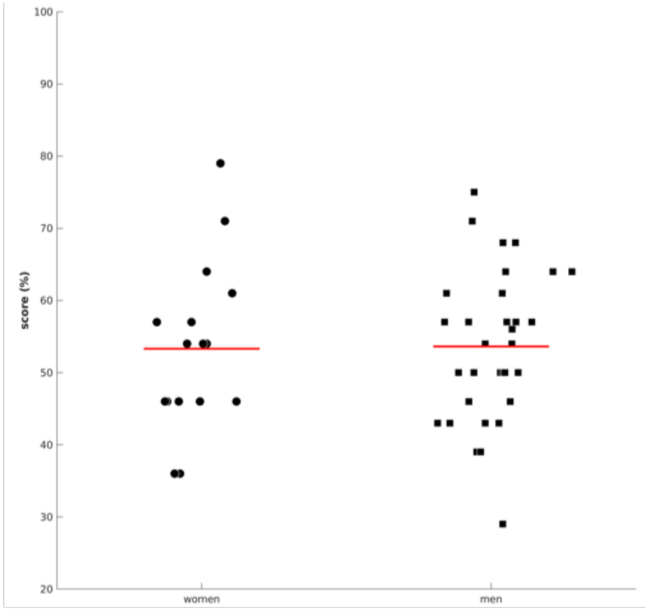


**Supplemental Figure 2 |** In order to assess whether variability in the results is driven by middle-aged participants, we assessed whether individuals above 35 years of age performed similarly to younger adults. No age-related differences were observed in either sex group (older women: 54.5±14.4%, young women: 52.9±11.3%, $t$=0.227, $p$=0.823; older men: 50.4±7.5%, young men: 54.5±11.0%, $t$=0.918, $p$=0.366). Thus, we combined data across age strata in each group and compared scores. We observed no sex differences in CSST-D performance (women: 53.3±11.7%; men: 53.6±10.4&; $t$=0.094, $p$=0.925).

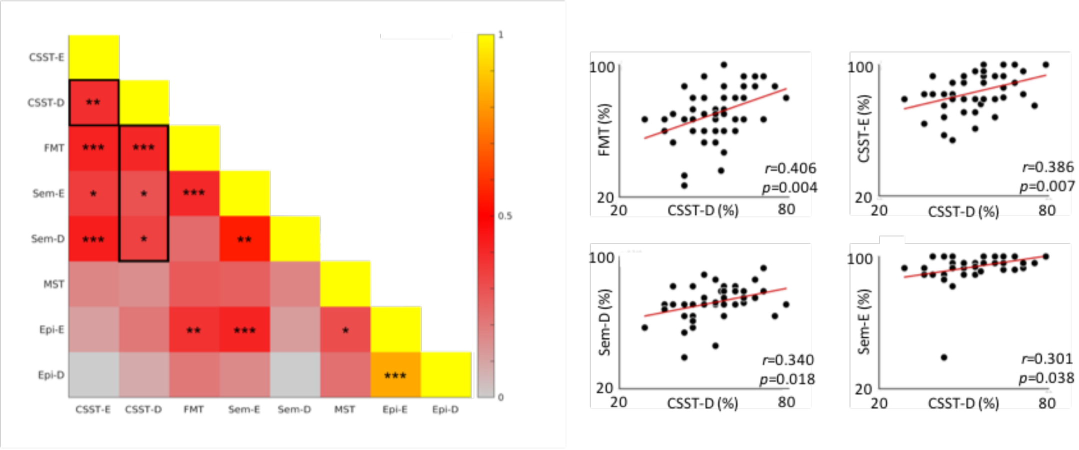


**Supplemental Figure 3 |** *left***:** correlation matrix of performance across all tasks, including easy conditions. Outlined area shows tasks with which CSST-D shows significant associations (*$p$<0.05; **$p$<0.01; ***$p$<0.005). *right***:** scatter plot of most significant associations with other tasks (FMT: Four Mountains Task; CSST-D/E: Conformational Shift Spatial Task-Difficult/Easy; Sem-D/E: Semantic Task-Difficult/Easy; Epi-D/E: Episodic Difficult/Easy; MST: Mnemonic Similarity/Discrimination Task)

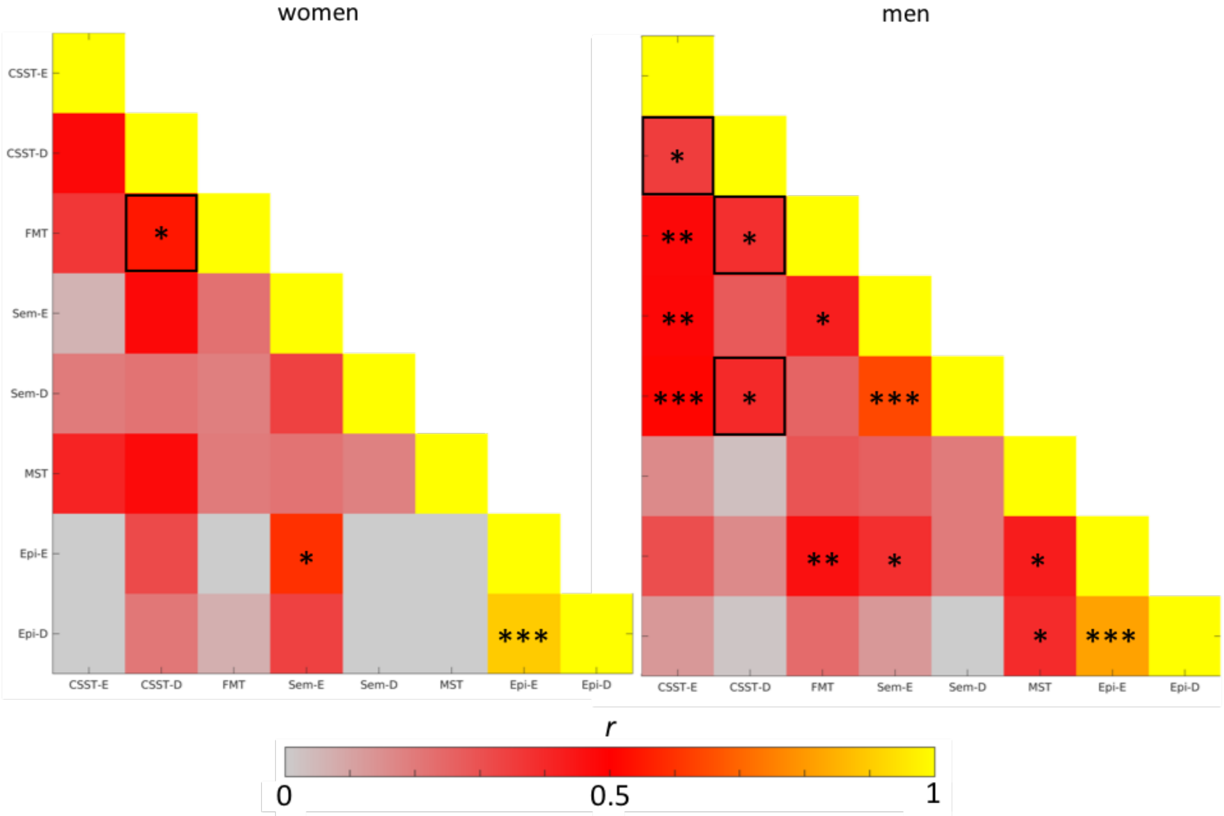


**Supplemental Figure 4 |** Correlation matrix of performance across all tasks for women and men. Outlined areas show tasks with which CSST-D shows significant associations (*$p$<0.05; **$p$<0.01; ***$p$<0.005). (FMT: Four Mountains Task; CSST-D/E: Conformational Shift Spatial Task-Difficult/Easy; Sem-D/E: Semantic Task-Difficult/Easy; Epi-D/E: Episodic Difficult/Easy; MST: Mnemonic Similarity Task)

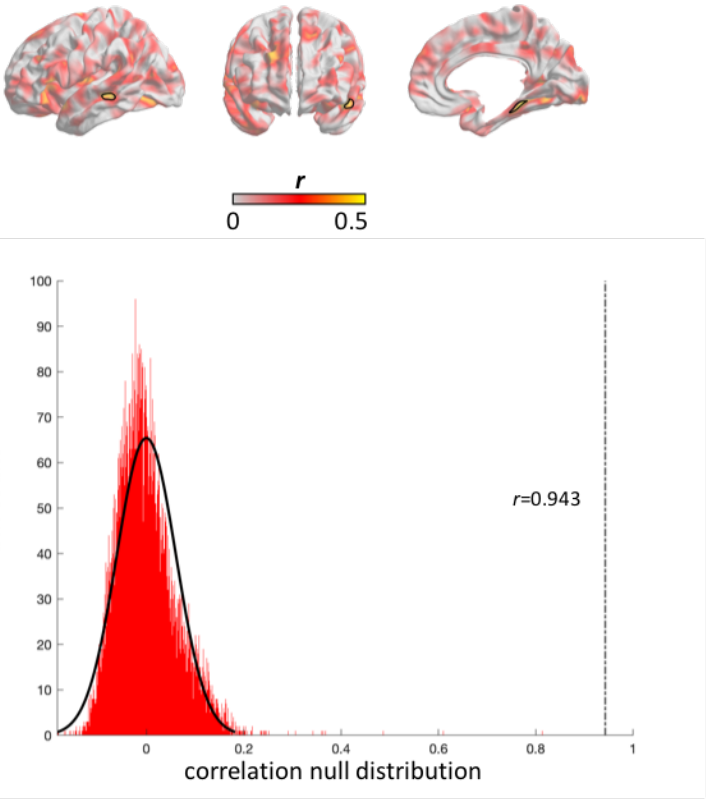


**Supplemental Figure 5 | top panel:** Pearson correlation coefficients of CSST-D performance on neocortical thickness after regressing out age and sex for right-handed participants (n=44). Highlighted clusters denote regions of significant association after multiple comparisons correction ($p_{FWE}<0.05$). **bottom panel:** a non-parametric null distribution was generated by correlating the *CSST-D x neocortical thickness* statistical *t* map with 10,000 permutated *t* maps of ***right-handed only*** *CSST-D x neocortical thickness*. Actual correlation between original maps is shown by the dashdotted line ($r$=0.943, non-parametric $p<0.001$).

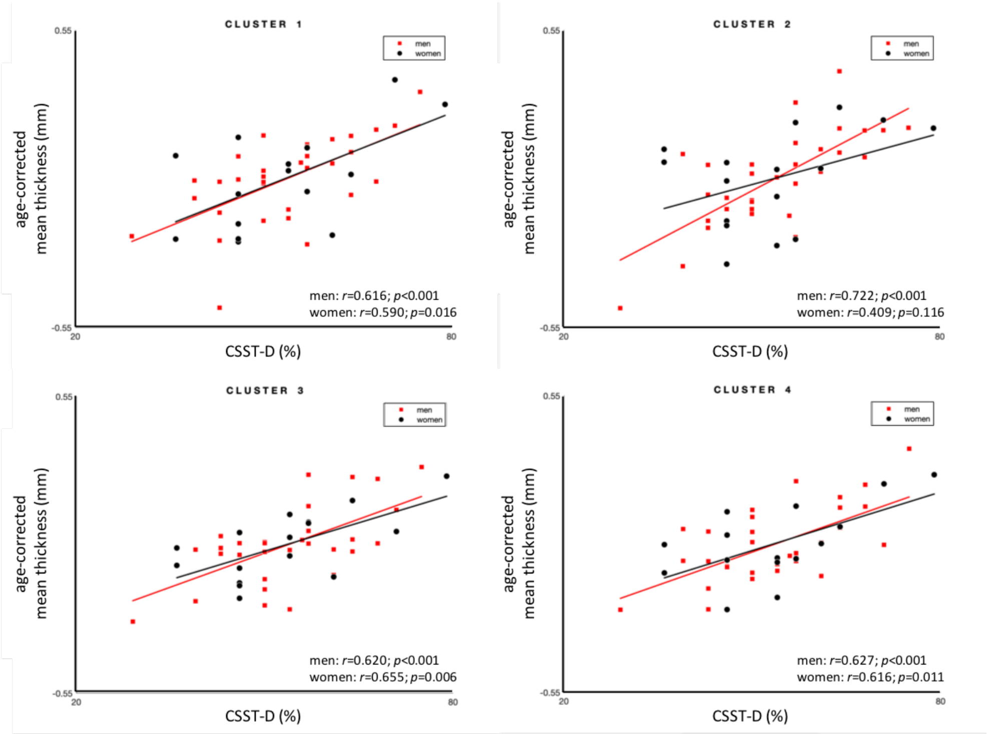


**Supplemental Figure 6 |** Controlling for age, we observed moderate-to-high associations between average cortical thickness and CSST-D scores for all clusters in men and women.

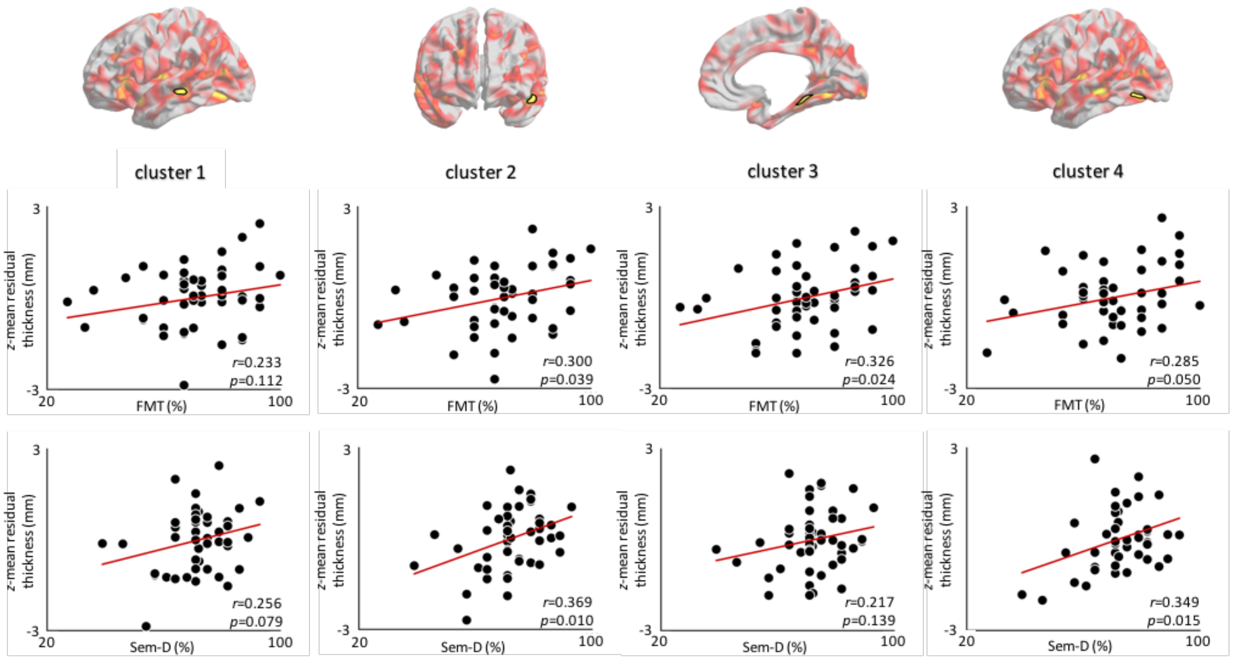


**Supplemental Figure 7 |** Cluster-wise associations between cortical thickness and scores for FMT (top row scatterplots) and Sem-D (bottom row scatterplots). Correlation coefficients ranged between r=0.233-0.326 for FMT (mean effect of 0.353) and between r=0.217-0.369 for Sem-D (mean effect of 0.373).

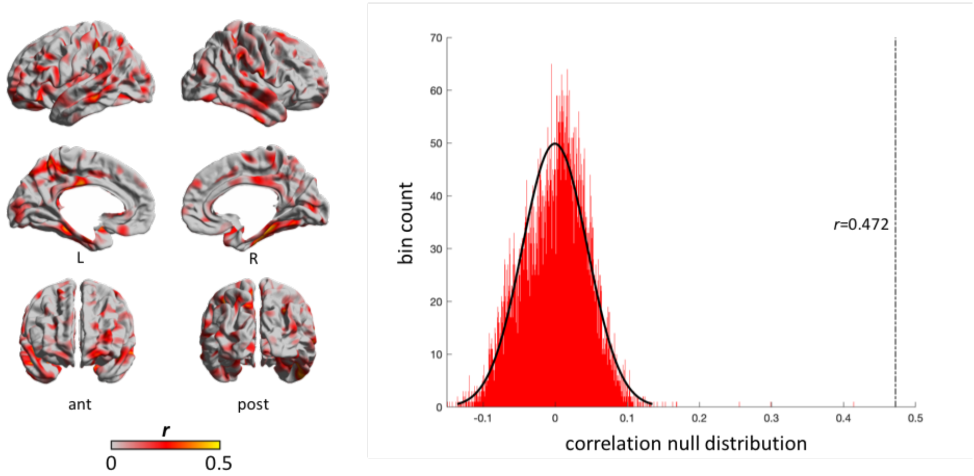


**Supplemental Figure 8 | left panel:** Pearson correlation coefficients of FMT performance on neocortical thickness after regressing out age and sex. **right panel:** a non-parametric null distribution was generated by correlating the *CSST-D x neocortical thickness* statistical *t* map with 10,000 permutated *t* maps of *FMT x neocortical thickness*. Actual correlation between original maps is shown by the dashdotted line ($r$=0.472, non-parametric $p$<0.001).

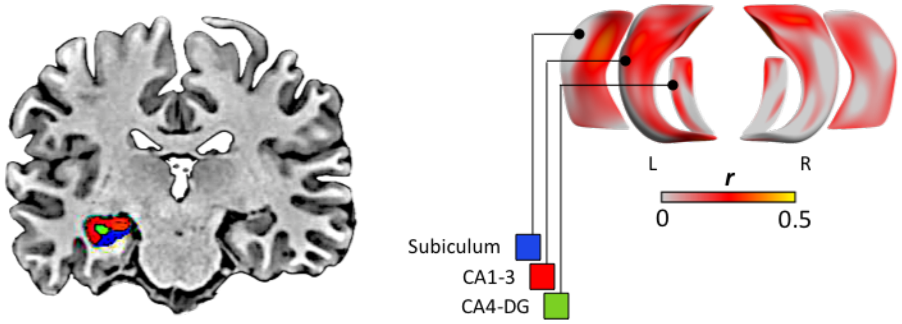


**Supplemental Figure 9 | left panel:** coronal section of the brain showing the hippocampal subfields.
**right panel:** uncorrected effect sizes shown on hippocampal subfield surfaces after regressing out age and sex.

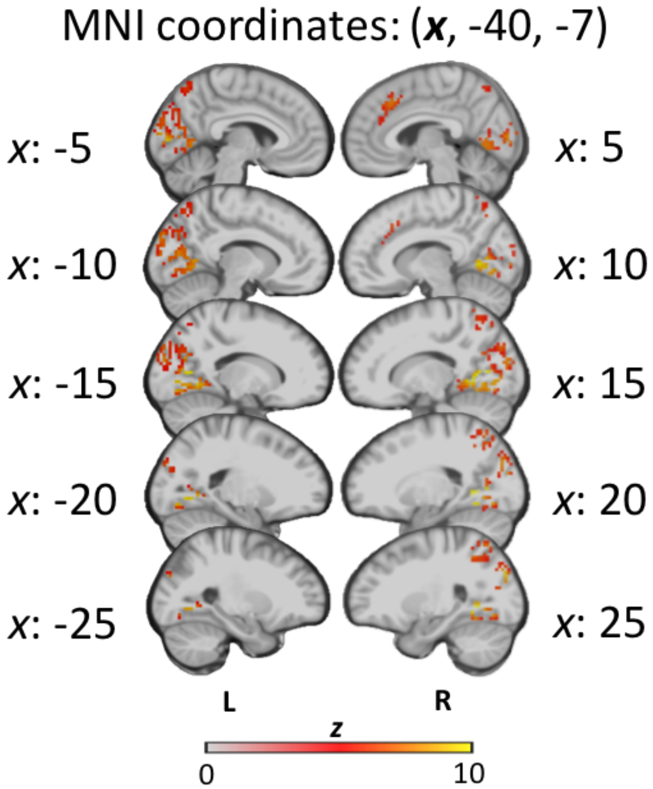


**Supplemental Figure 10 |** Group-level volumetric activation map for the contrast between retrieval and encoding.

| | | | | |
|---|---|---|---|---|
| FMT | 0.406 | | | |
| Sem-D | 0.340 | 0.237 | | |
| MST | 0.150 | 0.278 | 0.172 | |
| Epi-D | 0.083 | 0.206 | -0.058 | 0.224 |
| | CSST-D | FMT | Sem-D | MST |

**Supplemental Table1 |** Correlation coefficients of task performance scores (see **Figure 1b**)

| | successful | unsuccessful |
|---|---|---|
| CSST-E | 23 ± 3 (15-28) | 5 ± 3 (0-13) |
| CSST-D | 15 ± 3 (8-22) | 13 ± 3 (6-20) |
| Total | 38 ± 5 (28-50) | 18 ± 5 (6-28) |

**Supplemental Table 2 |** Number of successful and unsuccessful trials in each condition of the CSST reported as the mean ± SD (range)

| MNI x,y,z {mm} | peak T | peak p(unc) | peak p(FWE-corr) |
|---|---|---|---|
| 18 -64 5 | 14.87 | <0.001 | <0.001 |
| 21 -58 -1 | 14.52 | <0.001 | <0.001 |
| 12 -58 2 | 11.74 | <0.001 | <0.001 |
| -15 -67 8 | 11.24 | <0.001 | <0.001 |
| -18 -76 -4 | 11.16 | <0.001 | <0.001 |
| -21 -64 -4 | 10.7 | <0.001 | <0.001 |
| 33 23 2 | 9.68 | <0.001 | <0.001 |
| 39 20 -13 | 7.13 | <0.001 | 0.001 |
| -39 -22 59 | 9.54 | <0.001 | <0.001 |
| -39 -37 41 | 7.99 | <0.001 | <0.001 |
| -33 -16 65 | 7.97 | <0.001 | <0.001 |
| 36 -49 50 | 9.17 | <0.001 | <0.001 |
| 27 -52 47 | 9.01 | <0.001 | <0.001 |
| 24 -67 59 | 8.2 | <0.001 | <0.001 |
| 6 26 41 | 8.46 | <0.001 | <0.001 |
| 6 38 23 | 6.37 | <0.001 | 0.008 |
| 45 -28 47 | 8.19 | <0.001 | <0.001 |
| 42 -37 47 | 7.91 | <0.001 | <0.001 |
| 51 -19 44 | 7.19 | <0.001 | 0.001 |
| 45 32 23 | 8.05 | <0.001 | <0.001 |
| -9 -70 53 | 7.14 | <0.001 | 0.001 |
| -12 -76 47 | 6.52 | <0.001 | 0.005 |

**Supplemental Table 3 |** Group-level volumetric statistics for contrast between retrieval and encoding across pooled CSST-E and CSST-D trials.
